# High Capacity poly(2-oxazoline) formulation of TLR 7/8 agonist extends survival in a chemo-insensitive, metastatic model of Lung Adenocarcinoma

**DOI:** 10.1101/2019.12.12.874198

**Authors:** Natasha Vinod, Duhyeong Hwang, Salma H. Azam, Amanda E. D. Van Swearingen, Elizabeth Wayne, Sloane C. Fussell, Marina Sokolsky-Papkov, Chad V. Pecot, Alexander V. Kabanov

**Affiliations:** Center for Nanotechnology in Drug Delivery and Division of Pharmacoengineering and Molecular Pharmaceutics, Eshelman School of Pharmacy, University of North Carolina at Chapel Hill, NC 27599, U.S.A.; Joint UNC/NC State Department of Biomedical Engineering, University of North Carolina, Chapel Hill, NC, 27599□7575 USA; Lineberger Comprehensive Cancer Center, University of North Carolina at Chapel Hill, Chapel Hill, NC, 27599, USA; Department of Biology, Department of Chemistry, University of North Carolina at Chapel Hill, Chapel Hill, NC, 27599, USA; Division of Hematology & Oncology, University of North Carolina at Chapel Hill, Chapel Hill, NC, 27599, USA; Department of Medicine, University of North Carolina at Chapel Hill, Chapel Hill, NC, 27599, USA; Laboratory of Chemical Design of Bionanomaterials, Faculty of Chemistry, M.V. Lomonosov Moscow State University, Moscow, 119992, Russia

## Abstract

About 40% of the NSCLC patients have Stage IV cancer at the time of diagnosis. The only viable treatment options for metastatic disease are systemic chemotherapy and immunotherapy. Nonetheless, chemoresistance remains a major cause of chemotherapy failure. New immunotherapeutic modalities such as anti-PD1 checkpoint blockade have shown promise; however, response to such strategies is highly variable across patients. Here, we show that our novel poly(2-oxazoline) (POx) based nanomicellar formulation of Resiquimod, an imidazoquinoline TLR 7/8 agonist, had a superior tumor inhibitory effect in a metastatic model of lung adenocarcinoma, relative to anti-PD1 immune checkpoint blockade therapy as well as platinum-based chemotherapy, which is the mainstay of treatment for NSCLC. Investigation of the in vivo immune status following Resiquimod PM (POx micellar formulation of Resiquimod) treatment showed that Resiquimod-based stimulation of antigen-presenting cells in the tumor microenvironment resulted in the mobilization of anti-tumor CD8^+^ immune response. Our study demonstrates the promise of optimally delivered and nano-formulated Resiquimod as a new immunomodulating therapeutic strategy for the treatment of metastatic NSCLC.

## Introduction

Non-Small Cell Lung Cancer (NSCLC) is the most commonly diagnosed lung malignancy (constituting 80-85%) and accounts for the majority of cancer-related deaths worldwide (*1*). Post-surgical recurrence and metastasis is a principal cause of mortality in the vast majority of NSCLC cases (*2*). Genomic profiling of lung cancer has led to the identification of targetable mutations, paving the way for targeted therapies; however, the benefits of such interventions are transient due to chemotherapeutic resistance stemming from the heterogeneity of the tumor microenvironment (*1*). The development of systemic treatments that target the tumor microenvironment is thus crucial for managing advanced NSCLC.

FDA approval of immune checkpoint blockade therapy has reshaped the landscape of NSCLC treatment. Programmed death 1, more commonly known as PD1, is an immune checkpoint protein expressed on T cells to regulate self-tolerance by inhibiting immune attack against self-cells. Interaction of PD1 with its ligand (PDL1), commonly expressed among macrophages and myeloid cells, results in negative feedback generation, thereby deterring T cell response. Some cancers overexpress PDL1 to suppress anti-tumor response by T cells (*3*). The use of antibodies against PD1 has shown a favorable outcome in cancers with a high expression of PDL1 (*4*). However, only a minority of PDL1 positive NSCLC patients respond to anti-PD1 therapy due in part to intratumoral and temporal heterogeneity of pathologically regulated PDL1 expression, underscoring the role of the pathophysiological state of the tumor microenvironment in dictating the treatment response to anti-PD1 treatment (*5,6*).

Advances have been made in understanding the paradoxical role of immune cells in cancer. Signaling interactions between cancer cells and neighboring immune cells lead to the protumorigenic evolution of the latter, yielding cells that lack anti-tumor properties (*7*). For instance, a major proportion of the tumor-associated macrophages (TAMs) display an alternatively activated endotype, causing a shift in the Th1/Th2 cytokine balance towards a more Th2-like (anti-inflammatory) activity, rendering an immunosuppressive niche conducive to tumor growth (*8*). Additionally, TAMs can dampen the adaptive immune response by impeding the tumor infiltration of CD8+ cytotoxic T cells (*9*). Importantly, cancers that lack tumor-penetrating T lymphocytes (“cold tumors”) are reportedly refractory to immunotherapy (*10*). Thus, treatment strategies aimed at stimulating the T cell immune response are essential for a durable anti-tumor effect.

Toll-like receptor (TLR) agonists constitute a family of immune-stimulating agents that have demonstrated promising immune-enhancing effects in human and animal models of cancers (*11*). Expressed primarily on innate immune cells, TLRs are transmembrane proteins that recognize pathogen-associated molecular patterns (PAMPs), making it an indispensable part of the innate and adaptive immunity. Imiquimod (Aldara®; Graceway Pharmaceuticals) is currently the only clinically approved TLR 7 agonist, administered topically for the treatment of skin malignancies (*12*). Topical administration, however, isn’t feasible for cancers that are not accessible from the skin. Nonetheless, clinical studies involving systemic administration of TLR agonists at high doses have reported concerns regarding toxicity related to “cytokine storm” ensuing from systemic activation of TLR, resulting in flu-like symptoms (*13*). Thus, a delivery technique that allows for preferential targeting of tumor microenvironment while avoiding normal organs is warranted.

The present study investigates the immunotherapeutic potential of intravenously administered, Poly-(2-oxazoline) (POx) based nanomicellar formulation of Resiquimod (Resiquimod PM), a TLR 7/8 agonist chemically related to Imiquimod, in a clinically relevant mouse model of metastatic NSCLC. POx is an amphiphilic triblock-copolymer composed of one hydrophobic block of poly(2-butyl-2-oxazoline) (BuOx) flanked by two hydrophilic blocks of (2-methyl-2-oxazoline) (MeOx). POx micelles exhibit an exceptionally high solubilization capacity for water-insoluble drugs, in both singular and multi-drug combinations (*14,15*). We leveraged the characteristic sub-100 nm size of POx micelles to decrease the dose-limiting toxicity of the TLR agonist by passive targeting to the tumor. Furthermore, we also evaluated the anti-cancer efficacy of established frontline therapies for NSCLC, including immune checkpoint blockade therapy and platinum-based chemotherapy in combination with chemosensitizers, in the same model of NSCLC.

## Results

### Characterization of Poly-(2-oxazoline) formulations

#### Coformulation of chemosensitizers and anticancer agent

Cisplatin is a standard of care in advanced NSCLC (*16*). Nonetheless, the initial response to cisplatin is often short-lived due to the development of drug resistance *(17)*. We hypothesized that cancer cells could be sensitized to chemotherapy by the use of chemosensitizers, agents that harbor a potential for reversing drug resistance (*18*). Three different chemosensitizers were evaluated for co-formulation with an alkylated prodrug of cisplatin (C_6_CP). 1) AZD7762: a chemosensitizer that can inhibit the DNA repair activity by checkpoint kinases following treatment with DNA damaging agents such as Cisplatin and can thus improve the therapeutic margin of chemotherapy (*19*). 2) VE-822: an inhibitor of ATR (ataxia telangiectasia and Rad3-related), a DNA damage response pathway that is exploited by cancer cells as a rescue strategy on being subjected to DNA damage (*20*). 3) AZD8055: an inhibitor of the mammalian target of Rapamycin (mTOR) kinase. mTOR is a serine/threonine kinase involved in the regulation of cell growth and autophagy (*21*). Mutation in the mTOR pathway is common in NSCLC, making it a suitable choice of a chemosensitizer (*22*).

The hydrophobicity of the C_6_CP allowed easy *incorporation* of cisplatin within the hydrophobic micelle core. C_6_CP and chemosensitizer coloaded micelles were coformulated at different feeding ratios of the two drugs. Four ratios of C_6_CP: chemosensitizer (4:8, 6:6, 8:4, 9:3) were screened, keeping the polymer amount fixed. A maximum loading capacity of the micelle as high as 45% by wt. was obtained with the C_6_CP: AZD7762 coloaded micelles at a loading ratio of 2:6 (feeding ratio of 4:8). Micelle size varied with the feeding ratios (Table 1). C_6_CP: VE-822 coloaded micelles exhibited high loading efficiency and high loading capacity for all the ratios tested. The C_6_CP: AZD8055 coloaded micelles, too, exhibited a high loading capacity for all ratios except the feeding ratio of 4:8 (**Supplementary Table 1**). The micelles were a few hundred nanometers in size (Table 1).

**Table 1.**
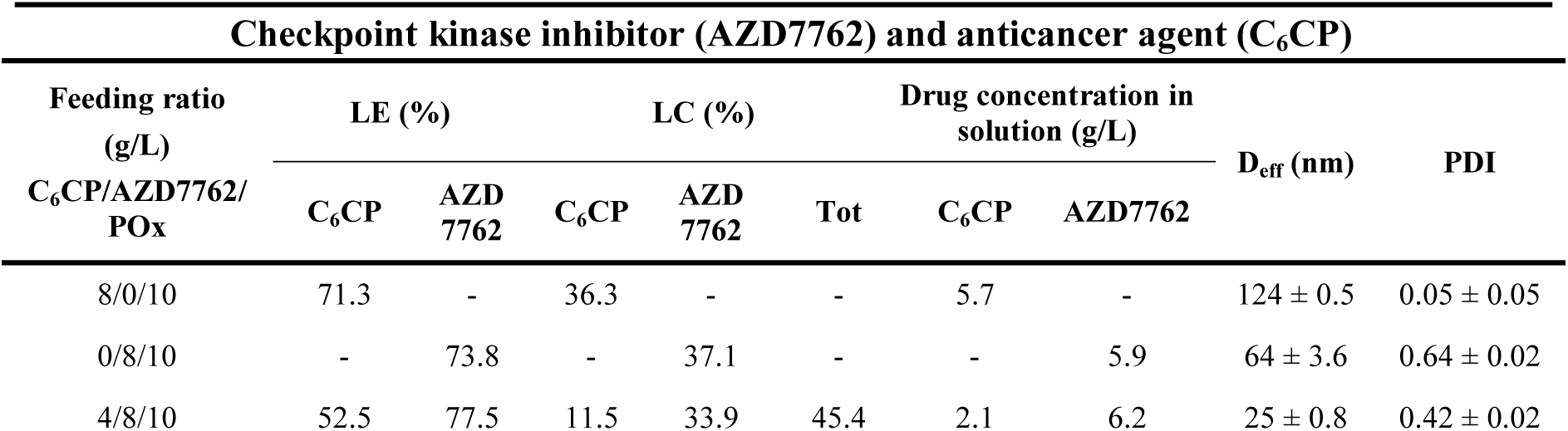
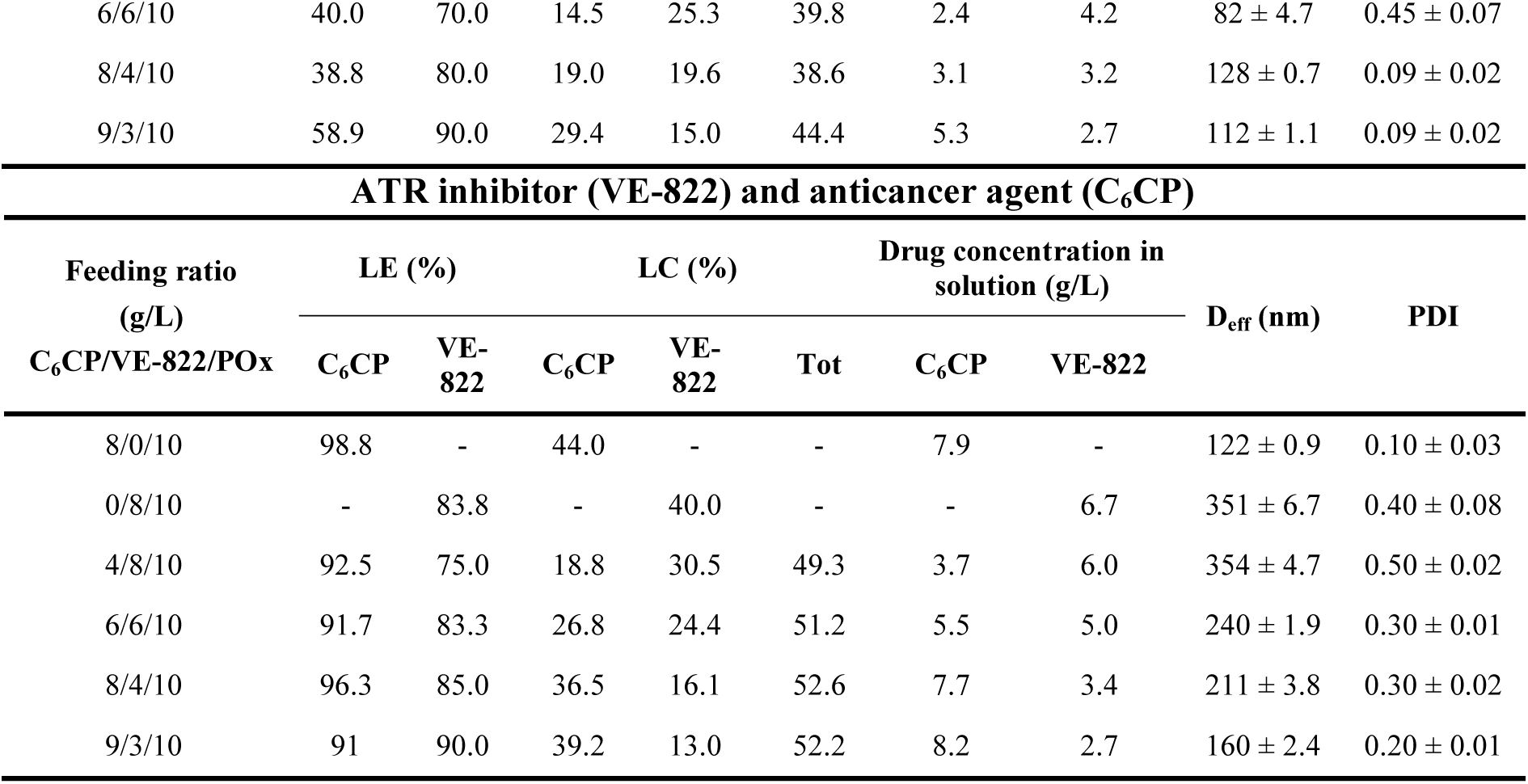
Characterization of POx micelles coloaded with anti-cancer agent and chemosensitizers.

The second chemotherapeutic drug screened for combination with chemosensitizer was Paclitaxel (PTX) since the POx micellar formulation of the drug has been extensively studied previously and has been found to exhibit a drug loading capacity superior to that of clinically approved Abraxane, resulting in a better anti-anticancer efficacy when administered at the maximum tolerated dose of the respective formulations (*14*). Four ratios of PTX: VE-822 were tested, and all the combination ratios exhibited high total loading capacity **Supplementary Table S1)**.

#### Resiquimod PM

Resiquimod is an imidazoquinoline immune response modifier (IRM)^12^. We sought to formulate resiquimod in POx micelles for safe intravenous administration in mice. By keeping the polymer amount constant and incrementally increasing the drug amount, different feeding ratios of POx/Resiquimod were examined. Resiquimod was well solubilized even at a high feeding ratio of 8/10, yielding a drug concentration of 7 mg/mL in saline and a loading capacity of 41% by wt. The size distribution obtained from DLS indicated the presence of small and monodisperse particles for feeding ratios 2/10 through 8/10 (Table 2, Figure 1a). This was corroborated by TEM, which showed small and spherical particles of about 20 nm in size (Figure 1b, 1c).

**Table 2.** Characterization of POx micelles loaded with TLR 7/8 agonist (Resiquimod)

| Feeding ratio<br>(g/L)<br>Resiquimod/POx | LE (%) | LC (%) | Drug concentration<br>in solution (g/L) | D<sub>eff</sub> (nm) | PDI |
| --- | --- | --- | --- | --- | --- |
| 2/10 | 90.0 | 15.0 | 1.8 | 24.6 ± 0.9 | 0.16 ± 0.01 |
| 4/10 | 97.5 | 28.0 | 3.9 | 24.5 ± 0.8 | 0.18 ± 0.02 |
| 8/10 | 87.5 | 41.0 | 7.0 | 25.9 ± 0.2 | 0.21 ± 0.01 |

**Fig. 1.**
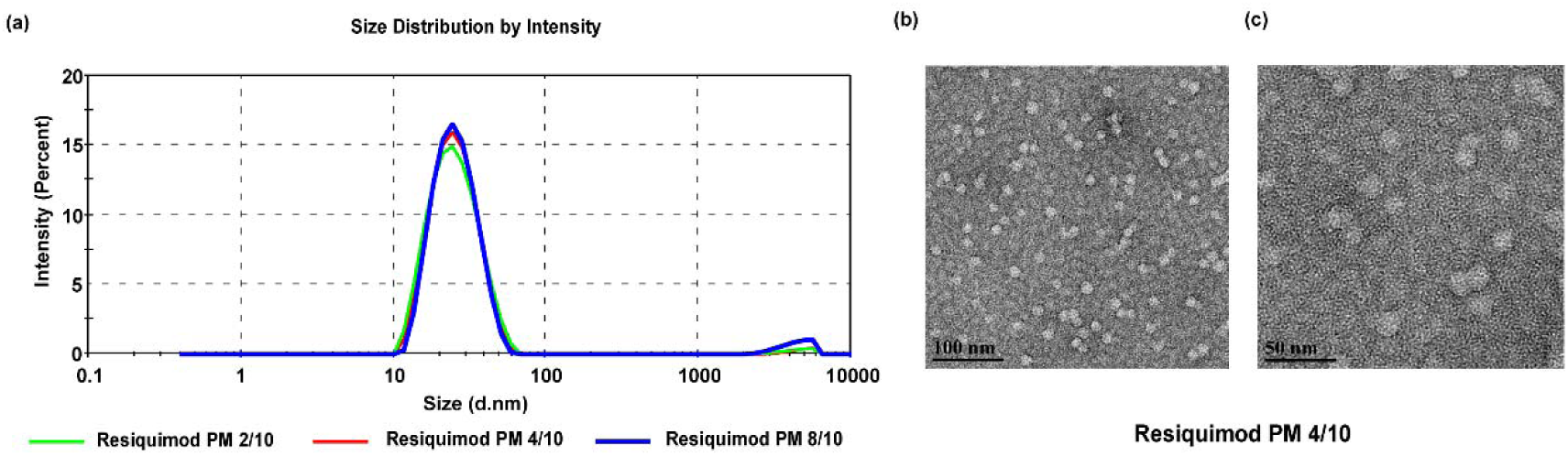
Physical characterization of Resiquimod PM. **(a)** Particle size distribution of Resiquimod PM at different feeding ratios as a function of intensity (percent), measured by dynamic light scattering; **(b, c)** Transmission electron micrographs at different magnifications show the spherical morphology of Resiquimod PM particles (4/10 g/L) and illustrate the uniformity of particle shape and size.

### In-vitro cytotoxicity

Anticancer agent (C_6_CP) and chemosensitizers (AZD7762, VE-822, AZD8055) alone and in combination were tested for their *in vitro* cytotoxicity against 344SQ lung adenocarcinoma cell line, with concentrations ranging from 0.256 ng/mL to 100 ug/mL. A dose-dependent decrease in cell viability was observed in all the treatments. The IC_50_ values were 0.01 ug/mL, 0.09 ug/mL, 0.4 mg/mL and 0.2 mg/mL for C_6_CP, AZD7762, VE-822 and AZD8055 respectively. The combinations showed a marked increase in cytotoxicity with IC_50_ values substantially lower than (by 5 folds for C_6_CP/AZD7762, 2 folds for C_6_CP/VE-822 and 2 folds for C_6_CP/AZD8055) either of the drugs alone, with the exception of C_6_CP (Figure 2, **Supplementary Figure 1a**). In contrast, the treatment of 344SQ cells with PTX produced a lower dose-dependent decrease in viability. Previous research has shown that increased exposure time in conjunction with increased dose improved the cytotoxicity of PTX (*23*); however, in the present study the cytotoxicity of PTX was found to be modest despite exposure for 72h. Nonetheless, the combination of PTX with VE-822 resulted in significantly higher cytotoxicity due to the chemosensitizing activity of VE-822 (**Supplementary Figure 1c**).

**Fig. 2.**
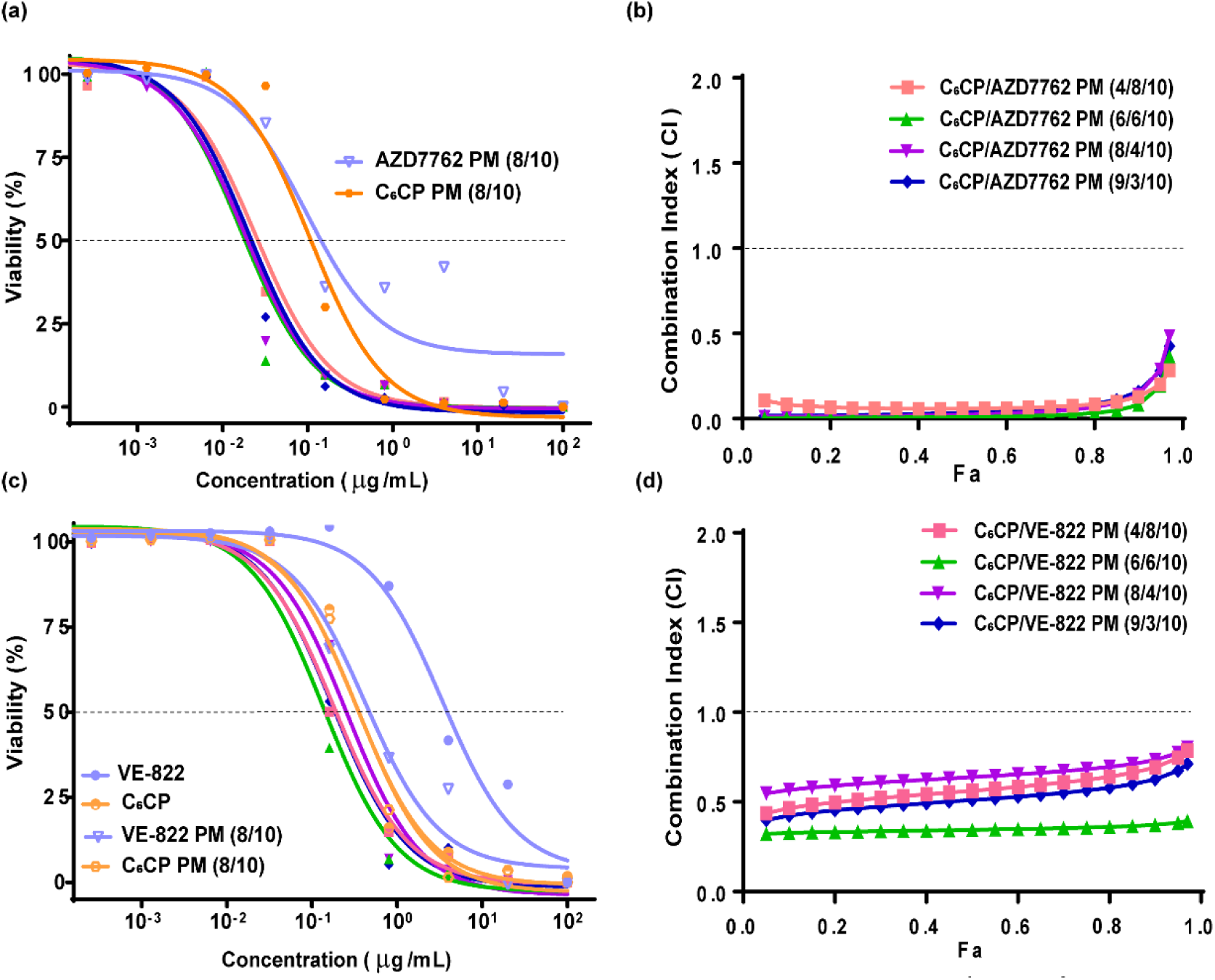
*In vitro* cytotoxicity of anticancer agent and chemosensitizers against 344SQ lung adenocarcinoma cell line. **(a, c)**. Dose-response curves of free and micelle incorporated drugs and drug combinations in 344SQ cell line after 72h of treatment. The data was fit into sigmoidal curve using non-linear regression. Data represent mean. n = 6. **(b, d)** Fa-CI plots of the C_6_CP/AZD7762 and C_6_CP/VE-822 combinations. Data represent mean. n = 6.

The drug synergy for different combination ratios was studied using the combination index (CI) theorem (isobologram equation) of Chou and Talalay, which states that a CI value of less than 1 represents synergism, whereas CI value greater than 1 indicates antagonism. Of note, the superadditive therapeutic effect of drug combinations is strongly influenced by the drug ratios *(15,24*). Interestingly, every feed ratio (4:8 through 9:3) of the C_6_CP/AZD7762 combination yielded CI < 0.3 for Fa ranging from 0.1 to 0.9, suggesting a strong synergy of the cotreatment. C_6_CP/VE-822 drug pair, too, depicted synergy for all feed ratios (CI < 1) with a pronounced synergy (CI < 0.5) for C_6_CP/VE-822 feed ratio of 4:8. C_6_CP/AZD8055 pair displayed maximum synergy at a feed ratio of 6:6, and while PTX/VE-822 showed synergistic effect at all feed ratios, maximum synergy was observed for the feed ratios of 8:4 and 9:3 (**Supplementary Figure 1b, and 1d**).

Due to the superior toxicity profile and synergistic effect of C_6_CP/AZD7762 and C_6_CP/VE-822 combinations, they were identified as lead candidates for *in vivo* study.

Resiquimod, on the other hand, did not exhibit any cytotoxic activity against 344SQ cells at concentrations ranging from 0.00128 ug/mL to 20 ug/mL (Figure 3). This observation is consistent with previous works that report Resiquimod as lacking a direct antineoplastic effect. However, its analog, imiquimod has been shown to exert a pro-apoptotic effect on a human skin cancer cell line. The disparate effects of the TLR agonists were speculated to be due to the differences in the subcellular localization of the two compounds (*25*).

**Fig. 3.**
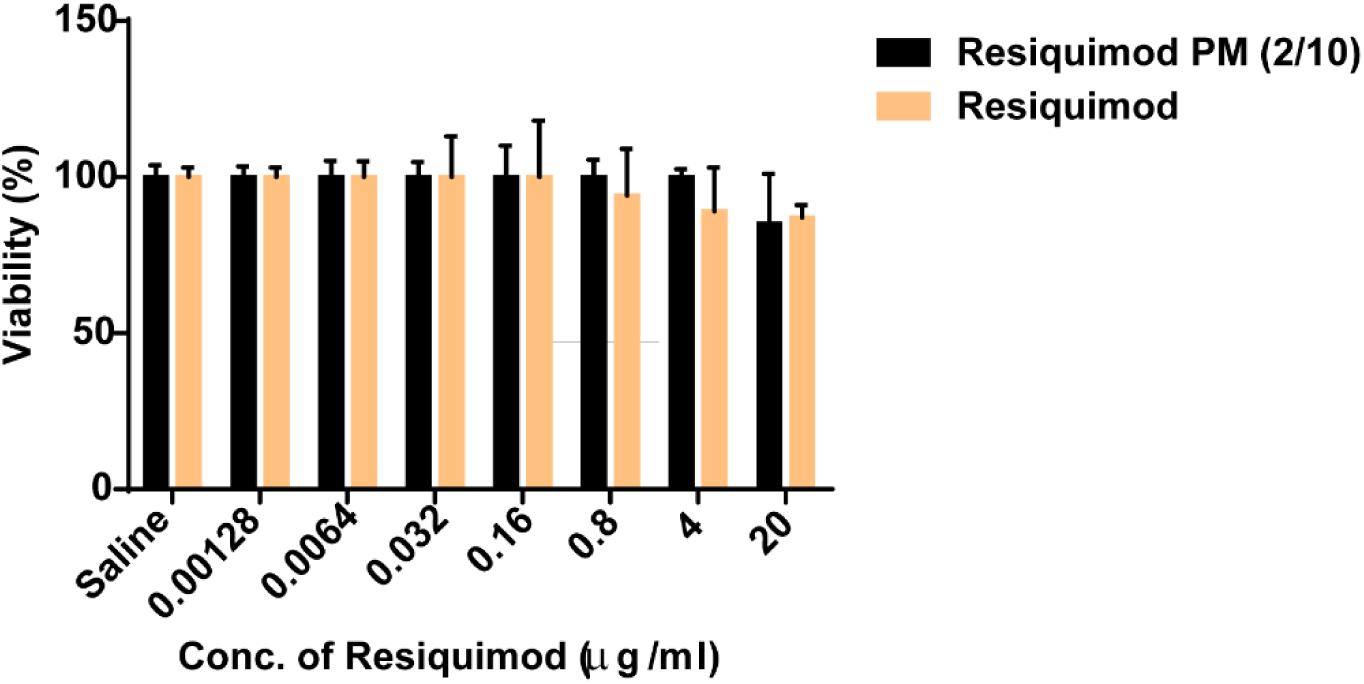
Cell viability of 344SQ lung adenocarcinoma cell line following 24h treatment with Resiquimod PM and free Resiquimod. Data represent mean ± SEM. n = 6.

### Characterization of *In vitro* Activation of BMDM by Resiquimod

Macrophages account for a major percentage of tumor-infiltrating leukocytes (*26*). Due to their plastic nature, TAMs are prime targets of cancer-mediated ‘reprogramming’ to a tolerogenic (Th1/Th2 low) phenotype. Immunotherapeutic strategies aimed at resetting the Th1/Th2 ratio to restore the tumoricidal activity of macrophages have shown promise in treating cancer (*27*). Accordingly, we sought to investigate the potential of Resiquimod PM to polarize murine bone marrow-derived macrophages to an anti-tumor phenotype (Th1/Th2 high) via TLR stimulation.

Conforming with a previous report, Resiquimod was found to lack cytotoxic effect on macrophages at the concentration used for the experiment (**Supplementary Figure 2**) (*28*). Resiquimod PM and free Resiquimod treatment of BMDM resulted in an increase in the mRNA expression of TNF-α, IL-1b, IL-6, and NOS2 (classical activation) in a manner similar to that of LPS, a TLR4 agonist (Figure 4a). Il-1b is an important Th-1 cytokine that primes anti-cancer immune response by the activation and expansion of CD4 and CD8 T effector cells (*29*). IL-6 signaling is again pivotal to the differentiation of T and B cells (*30*). The anti-tumor activity of macrophages ensues partly from NOS2 expression. NOS2 encodes inducible nitric oxide synthase (iNOS), an enzyme that catalyzes the production of tumoricidal reactive oxygen species (*8*). While the expression level of IL-6, IL-1b, and NOS2 for the Resiquimod and LPS treatment groups were considerably enhanced, TNF-α expression was relatively modest. This could be due to the 4h time-frame as the expression of TNF-α is reportedly low at early time points following macrophage stimulation (*31*).

**Fig. 4.**
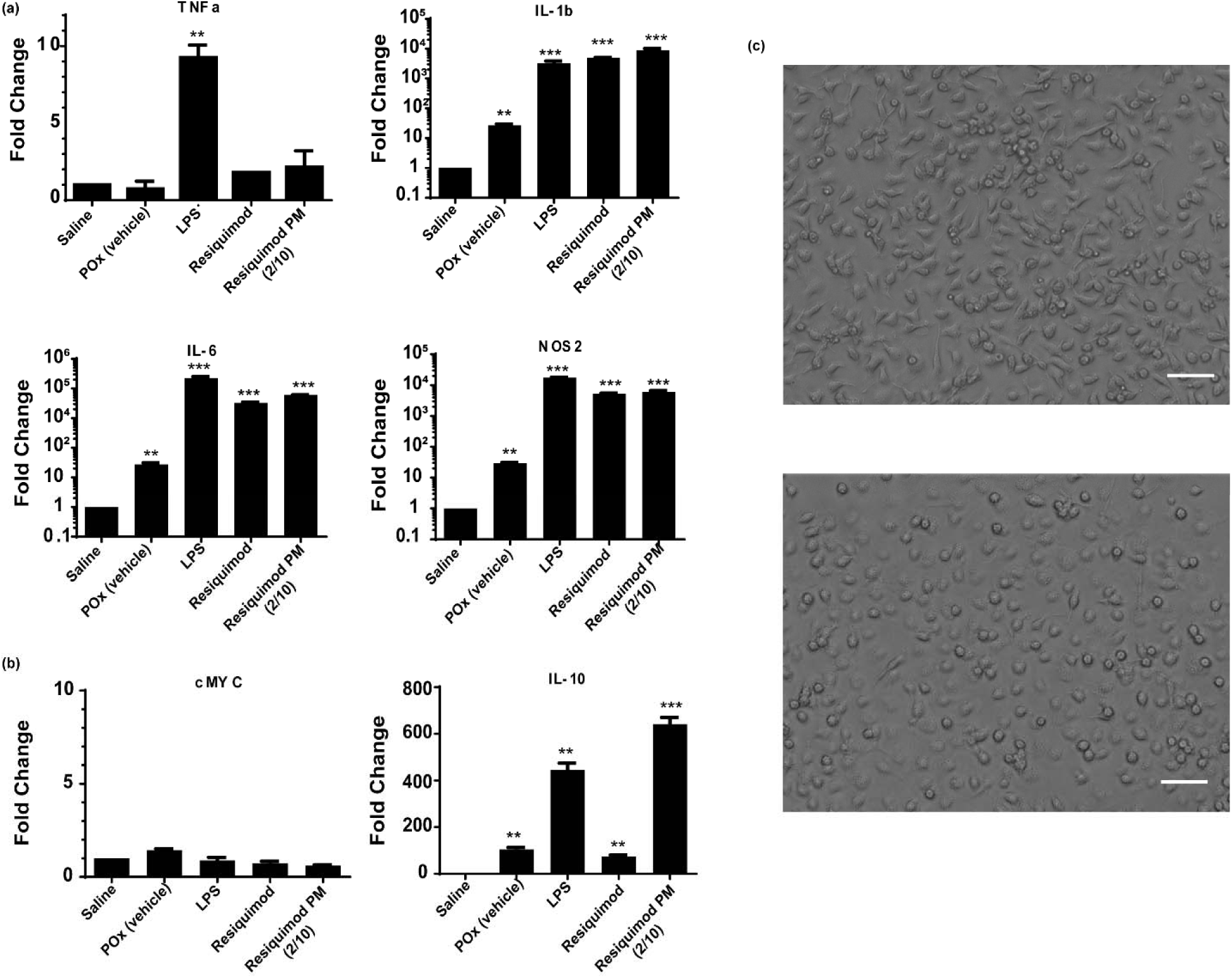
*In-vitro* activation of BMDM. **(a)** Relative mRNA expression of M1-like macrophage markers (TNFa, IL-1b, IL-6, NOS2) normalized to 18s **(b)** Relative mRNA expression of M2-like macrophage markers (cMYC, IL-10) normalized to 18s. Data represent mean ± SEM. n = 3. **p < 0.01, ***p < 0.001 computed by unpaired student t test with Welch’s correction. Significance level (α) was set at 0.05. **(c)** Cell morphology of resting macrophages (top) and M1 polarized macrophages following Resiquimod PM (2/10 g/L) treatment (bottom). Scale bar = 50 µm.

Analysis of alternatively activated macrophage markers showed a reciprocal downregulation of cMYC gene; however, Resiquimod and LPS treatment elicited increased expression of IL-10 (Figure 4b). This was not unexpected since IL-10 expression is known to counterbalance TNF-α production, resulting in a low TNF-α/IL-10 ratio. Increase in TNF-α expression over time is expected to thwart the production of IL-10.

Furthermore, POx (vehicle) was found to stimulate the expression of IL-1b, IL-6, and NOS2 in BMDM, albeit to a much lower extent than Resiquimod and LPS treatments (Figure 4a). This observation is in line with a study by Hou-Nan Wu et al. that looked at macrophage stimulation by amphiphilic polymers, where polymeric micelles induced the production of TNF-α and MCP-1 from macrophages in a time-dependent manner. However, following treatment of mice with these micelles, inflammatory mediators weren’t detected in the plasma of these animals (*32*). Therefore, we believe that the macrophage stimulation effect of POx micelles is not strong enough to warrant further investigation.

Following treatment with Resiquimod PM, cell morphology shifted from elongated structures in resting macrophages to round, and flattened structures, characteristic of Th1 activated macrophages (Figure 4c) (*33*).

### Estimation of Maximum Tolerated Dose

We have previously demonstrated the hematological and immunological safety of POx by assessment of liver and kidney function (blood chemistry panel), complement activation, and histopathology of major organs following repeated *i.v*. injections (q4d × 4) in mice (*14*). Therefore, no further toxicity analysis was conducted for the polymer alone in this study. Dose escalation study of single-agent POx micelles of C_6_CP, AZD7762, and VE-822 in 129/Sv mice served as a basis for identifying the doses for the combinations (data not shown). For both combinations, all the three tested doses (2.5/5, 5/10, 10/20 mg/kg and 10/10, 7.5/7.5, 5/5 mg/kg for C6CP/AZD7762 PM and C6CP/VE-822 PM respectively) were well tolerated by the mice. There was no incidence of death. Even for the highest tested dose, mice body weight didn’t fall below 5% of initial weight (Figure 5, **Supplementary Figures 3b, 3c**). Besides, no obvious behavioral abnormalities were observed in these mice. Accordingly, 10mg/kg of C_6_CP and 20mg/kg of AZD7762 for C_6_CP/AZD7762 and 10mg/kg of C_6_CP and 10mg/kg of VE-822 for C_6_CP/VE-822 were established as the MTD.

**Fig. 5.**
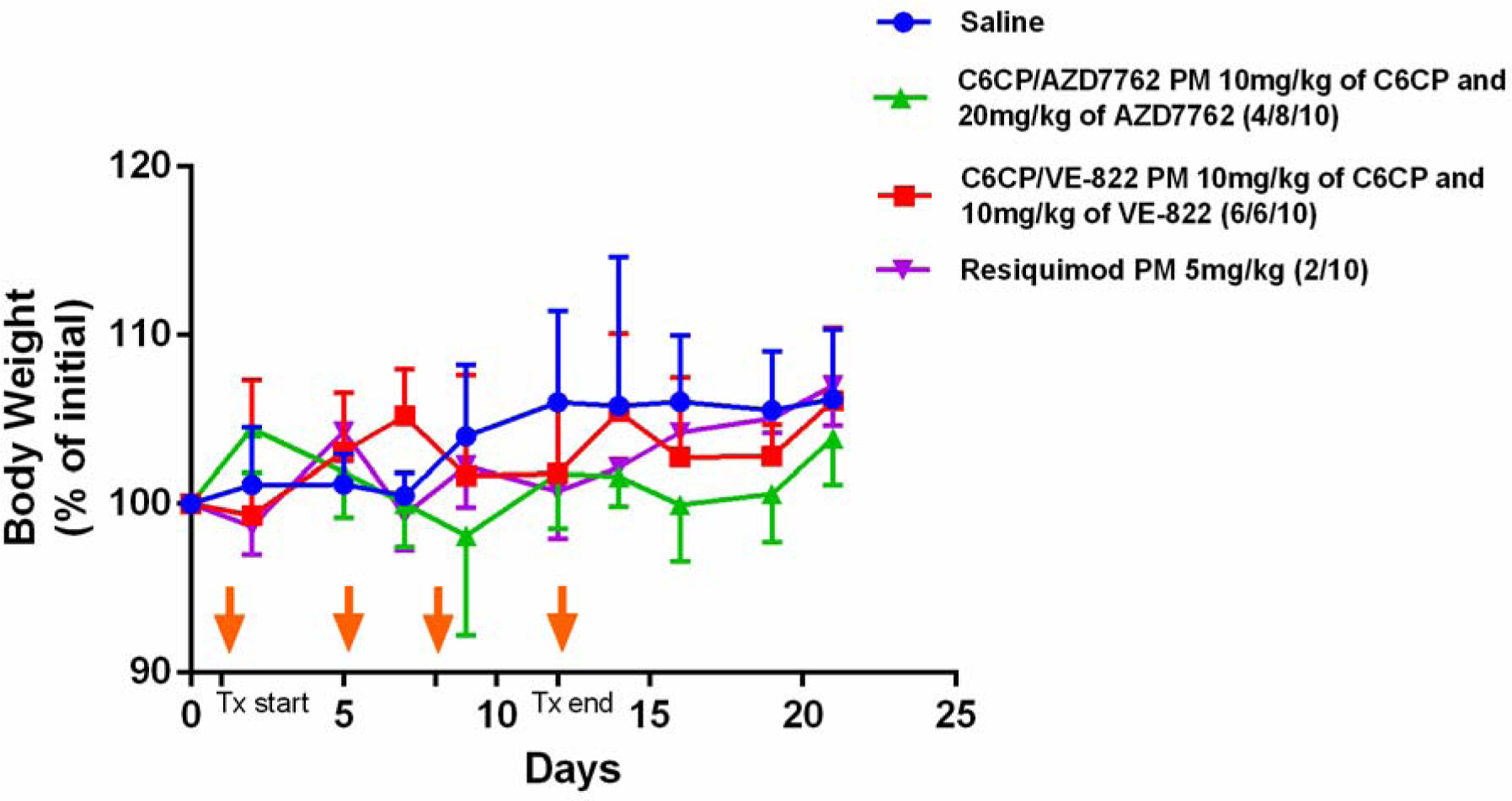
MTD study in healthy 129/Sv mice. Mice body weight (percent of initial) following four *i.v*. injections of POx formulations (q4d x 4). Data represent mean ± SEM. n = 3.

MTD finding studies for cancer immunotherapy is confounding since, unlike chemotherapy, higher doses do not necessarily increase efficacy. A non-linear dose-efficacy relationship of immune response modifiers makes it challenging to establish an MTD for these molecules. For this reason, most clinical studies involving immunomodulatory agents use doses that are below the MTD (*34*). As for Resiquimod PM, all the three tested doses were well-tolerated by the mice (Figure 5, **Supplementary Figure 3a**), as evidenced by the absence of any clinical signs. The body-weight changes of the Resiquimod PM treated group had a similar trend as the control group. Thus, 5mg/kg was identified as a safe dose for the in-vivo efficacy study.

### Tumor-inhibition study

We evaluated the anti-tumor efficacy of platinum-based chemotherapy when combined with chemosensitizers, and monotherapy with Resiquimod in an immune-competent, orthotopic model of lung adenocarcinoma (LUAD), prepared from 344SQ LUAD cell line derived from the metastasis of a genetically-engineered mouse model of LUAD carrying *Kras*^*G12D*^ and *p53*^*R172HΔG*^ mutations. The capacity to produce spontaneous metastases and thus recapitulate the pathophysiology of lung adenocarcinoma is a key strength of this model (*35*). Unexpectedly, neither of the combination drug PMs (C6CP/AZD7762 PM and C6CP/VE-822 PM) improved survival relative to the control group in spite of displaying a strong synergistic effect *in vitro*. The median survival for C_6_CP/AZD7762 PM was 24 days, and that for C_6_CP/VE-822 was 28 days, which was comparable to the untreated group. Furthermore, anti-PD1 treatment produced a modest improvement in survival, relative to the control (Figure 6a).

**Fig. 6.**
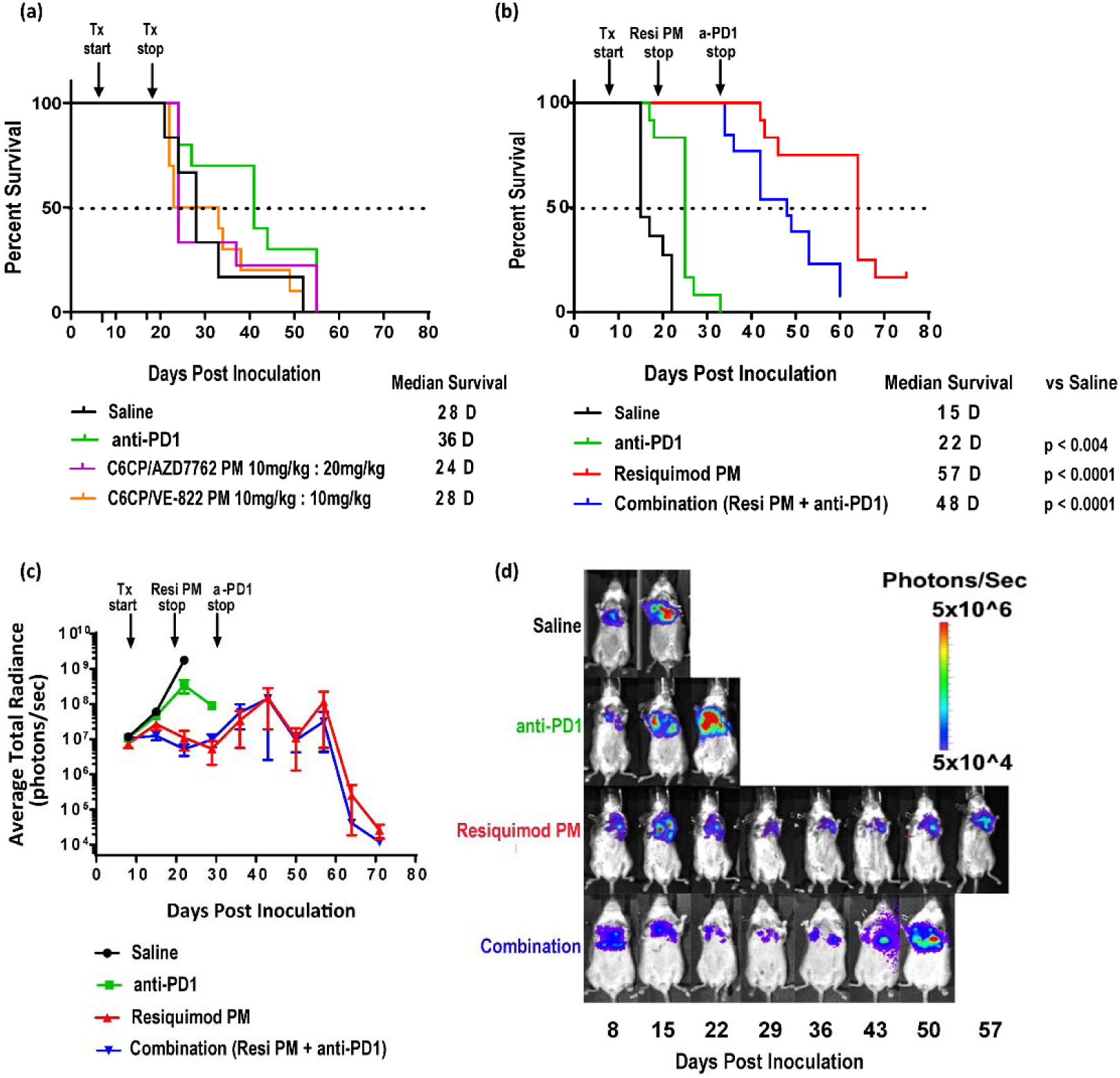
Tumor inhibition in 344SQ Lung Adenocarcinoma bearing mice. Kaplan-Meier survival plots of **(a)** tumor-bearing mice treated with four *i.v*. injections of Saline, C_6_CP/AZD7762 PM, and C_6_CP/VE-822 PM **(b)** tumor-bearing mice treated with four *i.v*. injections of Saline, Resiquimod PM, 8 *i.p*. injections of anti-PD1 antibody, and a combination of Resiquimod PM (4 *i.v*. injections) anti-PD1 antibody (8 *i.p. injections*), p values were computed by Log-rank (Mantel-cox) test. Significance level (α) was set at 0.05. (**c**) Quantification of BLI signal; data represent mean ± SEM. n = 13. (**d**) Representative IVIS images of mice from each treatment group on the days of the treatment.

On the contrary, Resiquimod PM monotherapy resulted in a pronounced increase in overall survival (Figure 6b). The median survival was 57 days for this group, which was a significant improvement considering the poor prognosis of this model of NSCLC. Luciferase expression of the 344SQ cell-line allowed for assessment of tumor growth by bioluminescence imaging. Despite lacking a direct anti-cancer effect (Figure 3), Resiquimod PM treatment substantially suppressed tumor progression (Figure 6c, 6d). Anti-PD1 monotherapy provided a modest benefit on tumor growth. Although the combination of PD1 with Resiquimod PM performed better than anti-PD1 alone, it didn’t provide any discernible benefit over Resiquimod PM monotherapy. A possible explanation for this is a lack of synergistic interaction between anti-PD1 and Resiquimod. Furthermore, the bodyweights of the mice from the Resiquimod PM and combination groups remained consistent when compared to mice from the saline and anti-PD1 groups (**Supplementary Figure 4**).

### Resiquimod controls LUAD growth by mediating host immune response

To uncover the immune-modulatory effect of Resiquimod cargo, the immune status of the tumor microenvironment was analyzed by flow cytometry at 48 hours after the second injection of Resiquimod PM in LUAD bearing mice. Given the vital role of macrophages in regulating the inflammatory response, we sought to examine its surface profile following TLR 7/8 stimulation. Tumors that received Resiquimod PM treatment showed an increased incidence of CD11b^+^/CD11c^-^/Ly6C^+^ monocytes (Figure 7). Ly6C^+^ monocytes are prone to differentiate into inflammatory macrophages and secrete Th1 cytokines that activate adaptive immune response (*36*). We next investigated the influence of Resiquimod treatment on dendritic cells. The CD11b^+^/CD11c^+^ expressing dendritic cell subset was found to be reduced in the treatment group in comparison to the control. This observation was consistent with the finding by Decker et al. that CD11c marker is downregulated upon activation of mouse dendritic cells by TLR stimulation (*37*). Accordingly, it is reasonable to conclude that dendritic cell activation by Resiquimod PM led to the downregulation of CD11c marker, yielding a low number of CD11c^+^ cells in the tumor. Finally, we examined if the stimulation of antigen-presenting cells led to the induction of T cell response. Flow analysis of cells triple stained for CD45^+^/CD3^+^/CD4^+^ and CD45^+^/CD3^+^/CD8^+^ revealed an increase in the CD8^+^ T cell population and an upward trend in CD4^+^ T cell population in the tumors of the treatment group, suggesting the ability of Resiquimod monotherapy to not only mount tumor-specific immune response by CD8^+^ T cells but also activate the CD4^+^ T cell population, required for the generation of memory immune response (*38*).

**Fig. 7.**
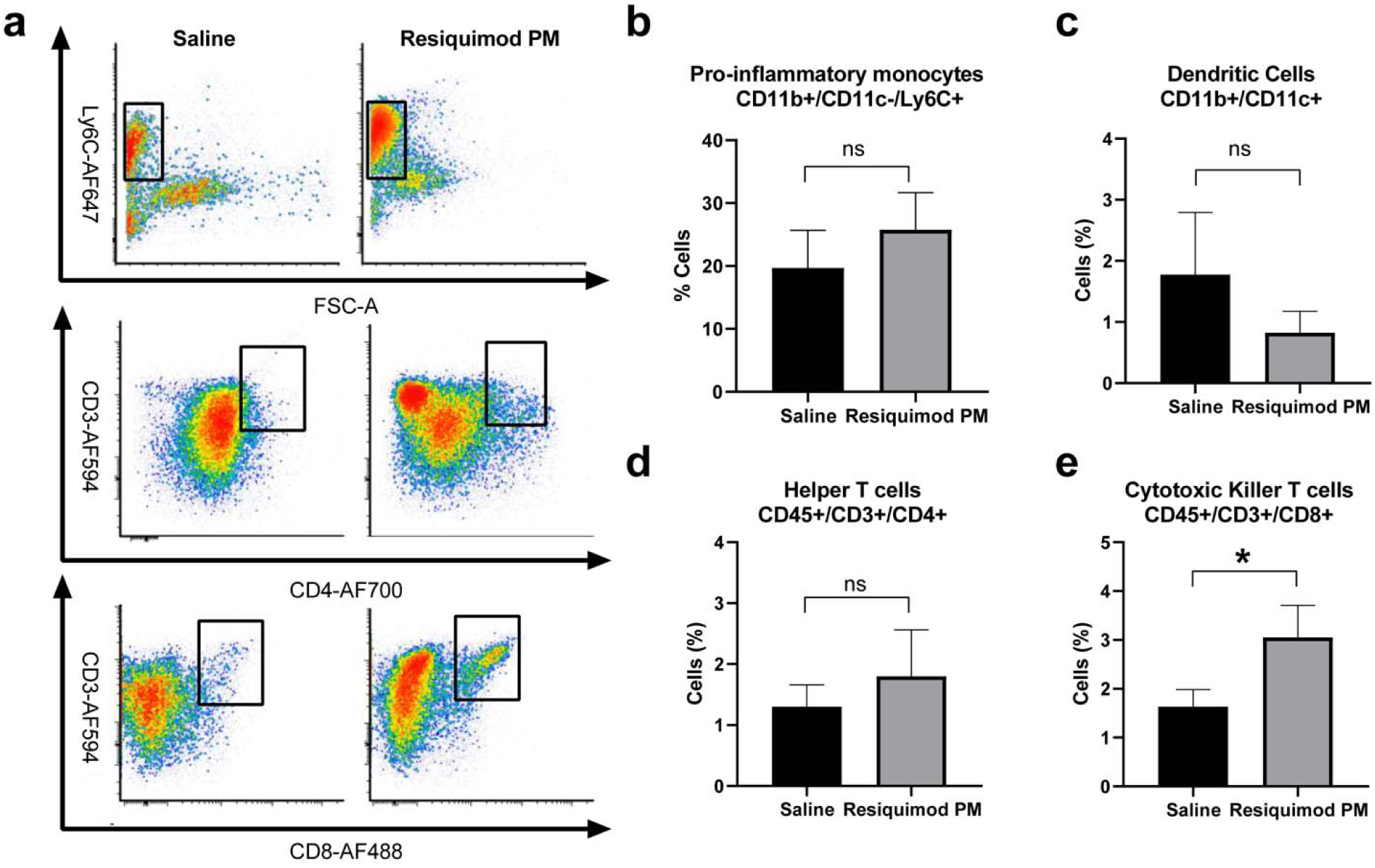
Resiquimod PM induces Th1 polarization of immune cells in the TME. **a)** Representative FACS plots of CD11b^+^/CD11c^-^/Ly6C^+^, CD45^+^/CD3^+^/CD4^+^ and CD45^+^/CD3^+^/CD8^+^ cell population from the tumors of mice treated with saline and Resiquimod PM. **b,c,d,e,)** Quantification of indicated population of cells. Data represent mean ± SEM. n = 4. *p < 0.05 computed by unpaired student t test with Welch’s correction. Significance level (α) was set at 0.05.

## Discussion

Our animal model of lung adenocarcinoma is developed by orthotopic injection of *Kras/p53* cell line (344SQ) into the lung of a syngeneic, immune-competent host. The 344SQ cell line is predisposed to metastasis due to loss of miR-200 family, a negative regulator of EMT, and thus, metastasis (*39*). Metastatic lung adenocarcinomas harboring *Kras/p53* mutations are associated with significantly lower numbers of tumor-infiltrating CD8^+^ T cells (*4*). Insufficient numbers of tumor-infiltrating T cells preclude response to immunotherapeutic strategies such as anti-PDL1 therapy that primarily act on pre-existing anti-cancer T cells (*40*), necessitating alternative approaches that can circumvent this problem.

Resiquimod is 100 times more potent (on a weight basis) as an immune response modifier than imiquimod. However, clinical trials involving topical Resiquimod have shown limited success owing to the poor systemic absorption (<1%) of local dose, resulting in suboptimal serum levels of Resiquimod (*12, 41*). Here, we report that our novel POx based nanomicellar formulation of Resiquimod provides an apposite platform for the systemic administration of the TLR agonist. Indeed, Resiquimod PM was not only well-tolerated by mice at *i.v*. dose of 5mg/kg, but also extended the overall survival in LUAD bearing mice, outperforming anti-PD1 therapy. In contrast, despite exhibiting an excellent *in vitro* synergistic anti-cancer effect, chemosensitizers and anticancer drugs co-formulated in POx micelles didn’t display a therapeutic effect in LUAD mice when compared to the control group, underscoring the insensitivity of LUAD model to chemotherapeutic strategies.

While lacking a direct anti-tumor effect, Resiquimod functions by orchestrating immune modulation of the tumor microenvironment resulting in the mobilization of the anti-tumor immune response (*42*). Immunogenicity of Resiquimod is conferred by its close resemblance to purine bases found in RNA, which are natural ligands of TLR (*12*). Since TLRs 7/8 are principally located intracellularly (*12*), encapsulation of Resiquimod in POx micelles is particularly beneficial for easy access to endosomally located TLR 7/8 following endocytic internalization of Resiquimod PM by immune cells. The association of Resiquimod and TLR 7/8 initiates the MyD88-dependent signaling cascade, which culminates in Th1 immune response. MyD88 (myeloid differentiation primary response 88) is an important adaptor protein that mediates the association between TLRs and IL-1R-associated kinases (IRAKs), and thereby triggers the activation of mitogen-activated protein kinases (MAPKs) and IκB kinase (IKK) complex, ultimately leading to the nuclear translocation and transcription of NFκB and subsequent induction of Th1 cytokines and chemokines (*12,43,44*). Th1 cytokine signaling potentiates the immune response against cancer and recruits more cells of the Th1-high endotype. Most notably, Th1 priming enhances the phagocytic activity of APCs and upregulates the expression of MHC and co-stimulatory molecules, resulting in the rapid phagocytosis of tumor cells and presentation of the tumor antigen to T cells in tumor-draining lymph node (TDLN), a prerequisite for the generation of tumor-specific immune response *(11)*. Our results indicate that Resiquimod PM can effectively polarize APCs (both macrophages and dendritic cells) to an anti-tumor phenotype *in-vivo* in LUAD bearing mice, corroborating our *in-vitro* study with BMDM, and concomitantly increase the infiltration of CD8^+^ T cells in the tumors, substantiating the potency of Resiquimod PM in generating a CTL response.

There is a growing understanding that the success of immunotherapy hinges on its ability to potentiate immune response to cancer by acting at the right location at the right time. While POx nanoformulation of Resiquimod addresses the former requisite, the latter can be addressed by using a dosing strategy that synergizes with the natural timing of immune response. From the recognition of tumor antigen to the infiltration of anti-tumor T cells, the development of immune response follows a coordinated sequence of events that takes several days (*45*). Accordingly, dosing schedule in immunotherapy should be optimized to allow enough time for maximum APC-T cell interaction. A limitation of this study is that we do not know if the dosing regimen chosen for the study is optimal, particularly for the Resiquimod PM and anti-PD1 combination, which didn’t show synergy. This will be investigated in the future with the help of suitable biomarkers that can offer a peek at the windows of opportunity for assessing the ideal time window for dosing to amplify the therapeutic efficacy of immunotherapy.

This study highlights the preeminent tumor-inhibition activity of Resiquimod PM brought about by effective immunomodulation of the tumor microenvironment and its potential to serve as an alternative to treatments that do not work on immunologically cold tumors. Although the investigation of the anti-tumor memory response was beyond the scope of this study, it is well recognized that activation and deployment of the adaptive immune surveillance generate long-term immunological memory that can counter cancer recurrence (*46*). Despite a promising therapeutic profile, a major bottleneck to clinical translation of Resiquimod is toxicity arising from systemic inflammation (cytokine storm). To this end, we have previously demonstrated the favorable pharmacokinetics of POx micelles in mice, allowing for tumor-specific accumulation of micelle cargo *(14*). In summary, we demonstrate that by using POx micellar platform for the intravenous delivery of Resiquimod, we are able to significantly decrease the dose-limiting toxicities and thus improve the therapeutic index of a potent drug such as Resiquimod, which had limited success in the clinical studies.

## Materials and Methods

Triblock copolymer of P[MeOx_35_-b-BuOx_34_-b-MeOx_35_]-piperazine (Mn=13kDa, Mw/Mn=1.14) was synthesized by living cationic ring-opening polymerization of 2-oxazolines as described previously (*47*). ^1^H NMR spectrum was obtained using Bruker Avance III 400 MHz spectrometer and analyzed using MestReNova (11.0) software. The molecular weight distribution of the polymer was measured by Gel Permeation Chromatography (GPC) on a Viscotek VE2001 solvent sampling module. The alkylated prodrug of cisplatin (C_6_CP) was synthesized as described previously (*48*). Resiquimod was purchased from ApexBio (#B1054) and Rat IgG2a, κ anti-mouse PD1, RMP1-14 clone was purchased from BioXCell (#BE0146). All other reagents were purchased from Sigma-Aldrich and Fisher Scientific.

### POx micelle preparation and characterization

POx micelles were prepared by the thin-film hydration method. The polymer and drugs (Resiquimod, C6 Cisplatin prodrug, Paclitaxel, AZD7762, VE-822, and AZD8055) were dissolved in a common solvent and subjected to mild heating (45 °C) accompanied by constant nitrogen flow for complete removal of solvent to form a dried thin film. The thin film was subsequently hydrated with saline at the optimal temperature (RT for Resiquimod PM and 55°C for C6CP/AZD7762, C6CP/VE-822, C6CP/AZD8005 and PTX/VE-822 PMs) to get drug-loaded micelles.

The drug amount incorporated in the micelles was measured by reversed-phase high-pressure liquid chromatography on an Agilent 1200 HPLC system equipped with Chemstation software, using a nucleosil C18, 5 μm particle size column (L × I.D. 25 cm × 4.6 mm). The UV chromatograms of drugs were obtained using isocratic elution mode with a mobile phase of ACN/Water 60/40 (v/v) & 0.1% trifluoroacetic acid, operated at a flow rate of 1 ml/min and a column temperature of 40°C. The micelle samples were diluted 50 times with the mobile phase, and an injection volume of 10ul was used for all the samples. The drug loading capacity and loading efficiency of the POx micelles were calculated as described previously (*14*).

The size distribution of POx micelles was determined using dynamic light scattering (DLS) technique on a Zetasizer Nano ZS (Malvern Instruments Ltd., UK). Every sample was diluted 10 times with normal saline to a final polymer concentration of 1 g/L, and the intensity weighted Z average size was recorded for 3 measurements of each sample at a detection angle of 173° and a temperature of 25°C. The POX micelles were further characterized by transmission electron microscopy. A high-resolution JEOL 2010F FasTEM-200kV with a Gatan CCD camera was used for image acquisition. Diluted solutions of POX micelles were dropped onto the TEM grid and allowed to dry and stained with 1% uranyl acetate for 2 mins before TEM imaging.

### Cell Study

In-vitro cytotoxicity of POx formulations on 344SQ lung adenocarcinoma cell line was assessed by studying the cell viability following treatment with various concentrations of free drugs and polymeric formulations of C6CP, PTX, AZD7762, VE-822, AZD8055, C6CP/AZD7762, C6CP/VE-822, C6CP/AZD8055, PTX/VE-822 and Resiquimod, prepared by serial dilution in full medium. 344SQ cell line was kindly provided by Dr. John Kurie (MD Anderson Cancer Center, Houston, TX). The cells were cultured in RPMI (Gibco™) medium supplemented with 10% fetal bovine serum (FBS) and 1% Penicillin-Streptomycin at 37°C with 5% CO_2_. 5000 cells/well were seeded in 96-well plates and allowed to attach for 24h before treatment. 72h following drug treatment, cell viability was measured by the Dojindo Cell Counting Kit-8 (CCK-8) using the manufacturer’s protocol. The half-maximal inhibitory concentration (IC50) was calculated using the GraphPad Prism 6 software. Quantification of the synergistic effect of drug combinations was done using CompuSyn software based on the combination index theorem of Chou and Talalay.

### In-vitro Activation of Bone Marrow-Derived Macrophages (BMDM)

BMDM were derived from the femur bone marrow of FVB/NJ mice per previously published protocol (*49*). Briefly, bone marrow cells were extracted from the bone marrow of 6-8 weeks old mice and subjected to red blood lysis by ACK lysing buffer. The resulting cell suspension was maintained in Dulbecco’s modified Eagle medium (DMEM) supplemented with 10% FBS, 1% Penicillin-Streptomycin, and 10 ng/ml recombinant murine M-CSF for 10 days. On Day 11, the medium was replaced with CSF free medium, and on the following day, the cells were treated with free, and micelle incorporated Resiquimod. For analysis of the in-vitro polarization status of macrophages, total RNA was harvested from BMDM 4 hours post-treatment per Qiagen RNA extraction protocol (QIAGEN). RNA was then reverse transcribed to cDNA using iScript Kit (Bio-Rad). Using cDNA as a template, the gene expression of the *Tnfa, il1b, il6, nos2, cmyc, il10*, and *mrc2* was measured by qPCR (relative to *18s*) on QuantStudio 6 Flex Real-time PCR system (Applied Biosystems).

### Estimation of Maximum Tolerated Dose (MTD)

A dose-escalation study was employed to identify the highest safe dose (MTD). Tumor-free female 8 weeks old 129/Sv mice were segregated into groups of three, with each group subjected to increasing doses of drugs. Resiquimod PM (1, 3 and 5 mg/kg), C6CP/AZD7762 PM (2.5/5, 5/10, 10/20 mg/kg) and C6CP/ VE-822 PM (10/10, 7.5/7.5, 5/5 mg/kg) and normal saline (control) were injected intravenously following q4d × 4 regimen. Every mouse was assigned a unique ID. Bodyweight loss of 15% or greater and other signs of toxicity such as hunched posture and rough coat were set as the study endpoints. The mice were monitored every other day until the end of the study.

### Animal Tumor Model of NSCLC

344SQ murine lung adenocarcinoma cells expressing firefly luciferase and green fluorescent protein (in 50 μL of 1:1 mix of HBSS and BD Matrigel) were injected into the left lung of 8 weeks old female 129/Sv mice via intrapulmonary injection as described previously (*50*). Briefly, mice anesthetized with ketamine + xylazine + acepromazine were laid in lateral decubitus position, and an incision was made between ribs 10 and 11 to visualize and access the lung. The cell suspension was directly injected into the lung parenchyma at the lateral dorsal axillary line, following which the incision was closed using surgical clips. The animals were monitored until full recovery.

### In-vivo efficacy study

#### Chemotherapy in conjunction with chemosensitizers

2.5×10^3^ 344SQ-GFP/fLuc cells were orthotopically injected in the left lung of 8 weeks old 129/Sv mice. Treatments were commenced a week after tumor inoculation. Baseline bioluminescence was measured using IVIS lumina optical imaging system prior to treatment administration. Mice randomized into groups of ten received i.v. injections of the following: 1) Normal saline; 2) 10/20 mg/kg of C6CP/AZD7762 PM 3) 10/10 mg/kg of C6CP/VE-822 PM; and i.p. injection of anti-PD1antibody (250 ug/mouse) using q4d × 4 regimen. Mouse survival and body weight changes were monitored every other day. Tumor load was measured weekly by bioluminescence imaging. Mice exhibiting signs of distress such as labored breathing, restricted mobility, ruffled fur, hunched posture, weight loss of greater than 15%, moribund state were euthanized by carbon dioxide intoxication followed by cervical dislocation.

#### Immunotherapy alone and in combination with immune checkpoint blockade

A week after tumor inoculation (5×10^3^ 344SQ-GFP/fLuc cells in 50 μL of 1:1 mix of HBSS and BD Matrigel), the animals (n=13) received the following injections: 1) Normal saline (i.v.; q4d × 4); 2) 5 mg/kg of Resiquimod PM (i.v.; q4d × 4); 3) 250 ug/mouse anti-PD1 (i.p.; q4d × 8); 4) 5 mg/kg Resiquimod PM (i.v.; q4d × 4) + 250 ug/mouse anti-PD1 (i.p.; q4d × 8).

### Evaluation of tumor microenvironment modulation by POx/Resiquimod

Subcutaneous 344SQ lung adenocarcinoma (10^5^ 344SQ-GFP/fLuc cells) tumors were formed in 8 weeks old 129/Sv mice. The mice were then randomly split into treatment arm (5 mg/kg Resiquimod PM; n=8) and control arm (Normal saline; n=8). Each group received two i.v. injections of the respective treatments on days 8 and 11 post tumor inoculation.

#### Flow Cytometry

For examining the immune status of the tumors post-treatment, the subcutaneous tumors were resected 48 hrs after the second treatment. The harvested tumors were subjected to enzyme treatment (Collagenase 2mg/mL in HBSS; Dispase 2.5U/mL in HBSS; DNase 1 mg/mL in PBS) for an hour at 37°C while shaking and digested into a single-cell suspension. The cell suspension was then passed through a 40µM cell strainer. After the removal of red blood cells by ACK lysis buffer, the cells were resuspended in FACS buffer (500 mL of 1X PBS w/o Ca^2+^ or Mg^2+^ + 2 mM EDTA + 2% FBS) and counted for downstream staining. 1×10^6^ live cells were stained with zombie violet live/dead stain (Biolegend) as per supplier’s recommendations, and excess live/dead stain was removed by washing cells twice and resuspending in 50 µl FACS buffer. Next, cells were incubated with 1µg anti-mouse CD16/CD32 (TruStain FcX™; Biolegend) on ice for 15 minutes. The cells were then mixed with 50 µl mixture of various fluorescently labeled monoclonal antibodies against murine cell surface markers **(Supplementary Table S2)** and incubated for 30 min on ice in the dark. Finally, cells were rinsed, resuspended in 300µL FACS buffer, and was immediately fluorescence-activated on LSRII (BD; FACSDiva 8.0.1 software) at the UNC Flow Cytometry Facility. Data was acquired with forward (FSC) and side (SSC) scatter on a linear scale, while fluorescent signals were collected on a 5-decade log scale with a minimum of 100,000 events per sample. Non-stained harvested cells were used as the universal negative control. Compensation beads (ThermoFisher) were used for single-color control samples. Harvested spleen cells were used as positive controls for immune cell staining. Analysis of flow cytometry data was performed using FCS Express (DeNovo Software). All antibodies were purchased from Biolegend.

## Acknowledgments

The authors thank Charlene Santos, Mark Ross and Alain Valdivia from ASC for helping with *i.v./i.p*. injections. The flow analysis was performed at the UNC flow cytometry facility, which is supported in part by P30 CA016086 Cancer Center Core Support Grant to the UNC Lineberger Comprehensive Cancer Center. Transmission electron microscopy was performed by Amar Shankar Kumbhar at the Chapel Hill Analytical and Nanofabrication Laboratory, CHANL, a member of the North Carolina Research Triangle Nanotechnology Network, RTNN, which is supported by the National Science Foundation, Grant ECCS-1542015, as part of the National Nanotechnology Coordinated Infrastructure (NNCI).

## Author Contributions

Conceptualization: N.V., D.H., A.V.K. C.V.P., and M.S.P.; Methodology: C.V.P., S.H.A., A.E.D.V.S., N.V., D.H. and E.W.; Investigation: N.V., D.H., S.C.F., and E.W.; Formal Analysis: S.H.A., A.E.D.V.S., C.V.P., N.V. and E.W.; Funding acquisition: A.V.K. and C.V.K.; Writing – original draft preparation: N.V.; Writing – review and editing: C.V.P. and A.V.K.; Supervision: C.V.P., A.V.K. and M.S.P.

