## Supplementary material for "High Capacity poly(2-oxazoline) formulation of TLR 7/8 agonist extends survival in a chemo-insensitive, metastatic model of Lung Adenocarcinoma": Table S1, Fig. S1, Fig. S2, Fig. S3, Fig. S4, Table S2

**Author Affiliations:**

**Table S1 | Characterization of POx micelles coloaded with anti-cancer agent and chemosensitizers**

**mTOR kinase inhibitor (AZD8055) and anticancer** **agent (C_6_CP)**

| **Feeding ratio**  **(g/L)**  **C_6_CP/AZD8055/POx** | | **LE (%)** | | **LC (%)** | | | **Drug concentration in solution (g/L)** | | |
| --- | --- | --- | --- | --- | --- | --- | --- | --- | --- |
|  |  | **C_6_CP** | **AZD8055** | **C_6_CP** | **AZD8055** | **Tot** | | **C_6_CP** | **AZD8055** |
| 8/0/10 | 70.6 | | - | 36.1 | - | - | | 5.7 | - |
| 0/8/10 | - | | 43.4 | - | 25.8 | - | | - | 3.5 |
| 4/8/10 | 22.0 | | 39.13 | 6.28 | 22.3 | 28.6 | | 0.9 | 3.1 |
| 6/6/10 | 83.2 | | 91.2 | 24.38 | 26.7 | 51.1 | | 5.0 | 5.5 |
| 8/4/10 | 85.3 | | 91.0 | 33.33 | 17.8 | 51.1 | | 6.8 | 3.7 |
| 9/3/10 | 80.2 | | 102.0 | 35.6 | 15.1 | 50.7 | | 7.2 | 3.1 |

**ATR inhibitor (VE-822) and anticancer** **agent (PTX)**

| **Feeding ratio**  **(g/L)**  **PTX/VE-822/POx** | | **LE (%)** | | **LC (%)** | | | **Drug concentration in solution (g/L)** | | |
| --- | --- | --- | --- | --- | --- | --- | --- | --- | --- |
|  |  | **PTX** | **VE-822** | **PTX** | **VE-822** | **Tot** | | **PTX** | **VE-822** |
| 8/0/10 | 86.9 | | - | 41.0 | - | - | | 7.0 | - |
| 0/8/10 | - | | 83.3 | - | 40.0 | - | | - | 6.7 |
| 4/8/10 | 83.8 | | 77.8 | 17.1 | 31.9 | 48.9 | | 3.4 | 6.2 |
| 6/6/10 | 102.0 | | 91.5 | 28.4 | 25.4 | 53.7 | | 6.1 | 5.5 |
| 8/4/10 | 97.0 | | 89.3 | 36.4 | 16.7 | 53.1 | | 7.8 | 3.6 |
| 9/3/10 | 95.2 | | 86.7 | 40.5 | 12.3 | 52.8 | | 8.6 | 2.6 |

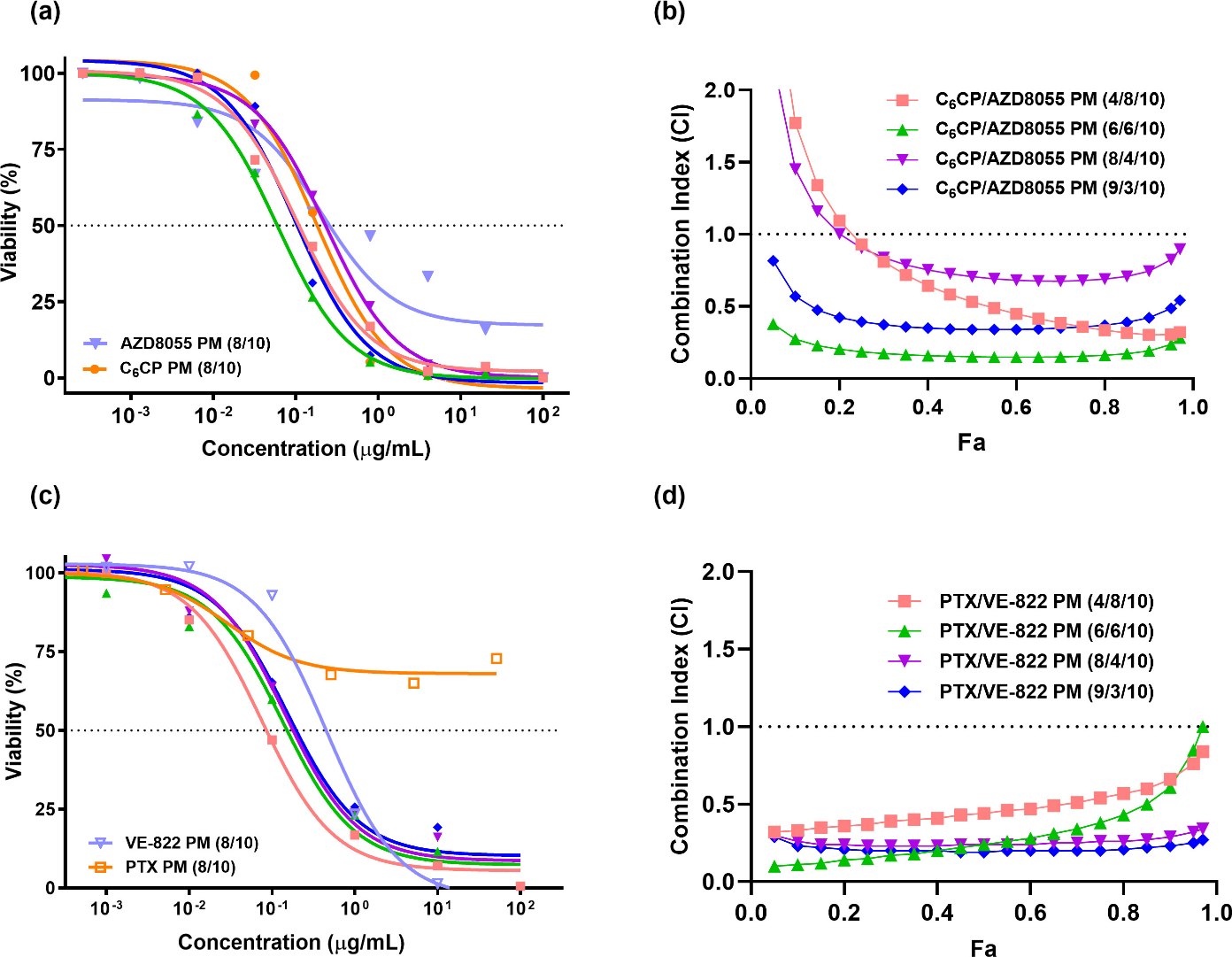

**Fig. S1. *In vitro* cytotoxicity of anticancer agent and chemosensitizers against 344SQ lung adenocarcinoma cell line (a,c)** Dose-response curves of free and micelle incorporated drugs and drug combinations in 344SQ cell line after 72h of treatment. The data was fit into sigmodal curve using non-linear regression. Data represent mean. n=6. **(b,d)** Fa-CI plots of the C_6_CP/AZD8055 and PTX/ VE-822 combinations. Data represent mean. n = 6.

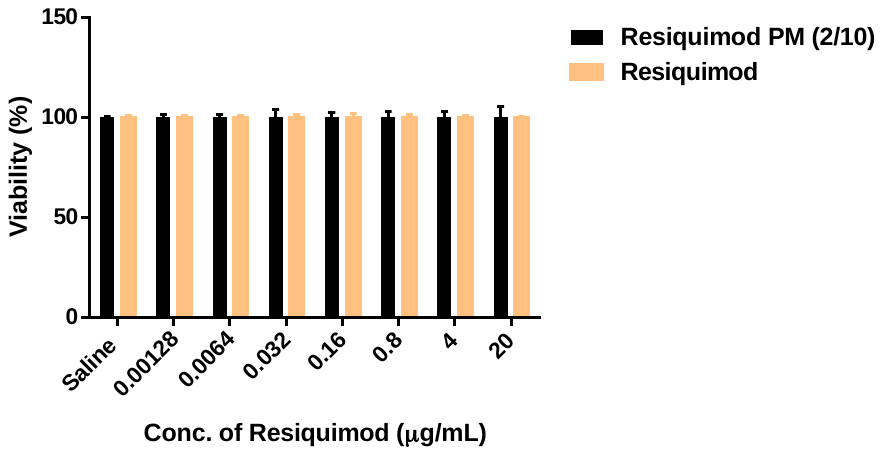

**Fig. S2.** Cell viability of BMDM following 24h treatment with Resiquimod PM and free Resiquimod. Data represent mean ± SEM. n = 6.

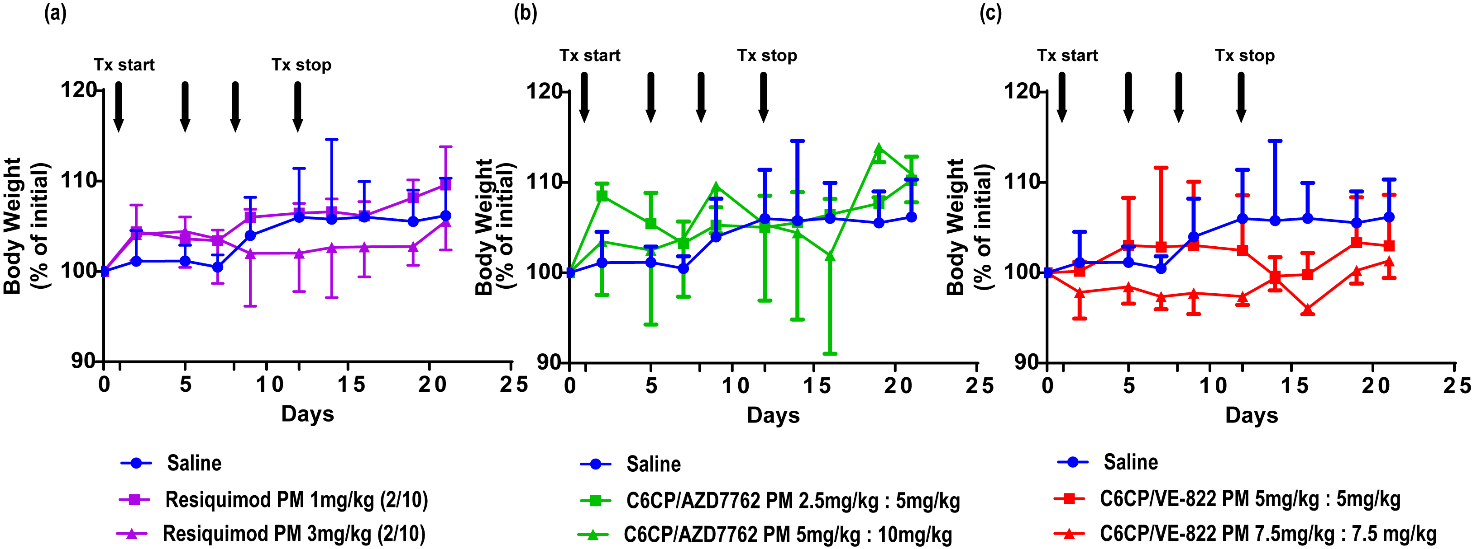

**Fig. S3. MTD study in healthy 129/Sv mice** Mice body weight (percent of initial) following four injections of **a)** Resiquimod PM (2/10 g/L) **b)** C_6_CP/AZD7762 PM **c)** C_6_CP/VE-822 PM (q4d x 4). A common control (saline) was used for all the groups. Data represent mean ± SEM. n = 3.

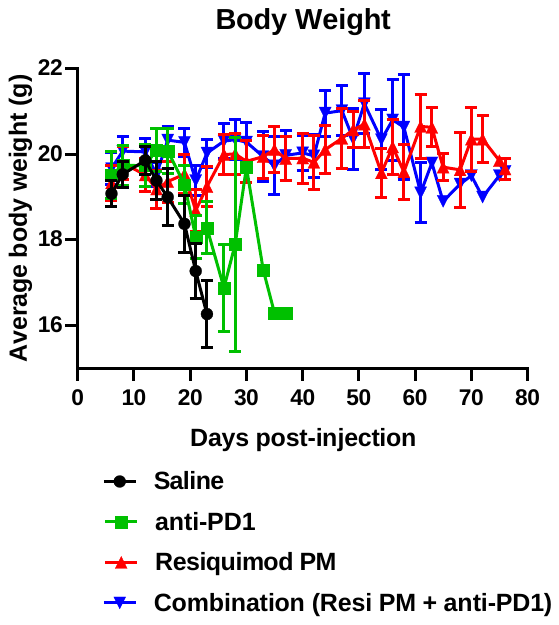

**Fig. S4.** Changes in the bodyweight of animals subjected to different treatments. Data represent mean ± SEM. n = 3.

**Table S2 | Description of Antibody-Fluorochrome pairs**

|  | **Myeloid Panel** | | **Lymphoid Panel** | |
| --- | --- | --- | --- | --- |
| **Antibody** | | **Fluorophore** | **Antibody** | **Fluorophore** |
| Ly6C | | AF647 | CD45 | PE |
| CD11b | | BV510 | CD8 | AF488 |
| CD11c | | APC/Cy7 | CD3 | AF594 |
| Ly6G | | AF594 | AF700 | AF700 |
